# Stop and go: exploring alternative mechanisms for task allocation in social insects - response and satisfaction thresholds trade off cost, accuracy, and speed differently

**DOI:** 10.1101/2024.05.13.593812

**Authors:** CM Lynch, RC Wilson, A Dornhaus

**Author notes:** Corresponding author (CL).

## Abstract

Division of labor, a key feature of many complex systems, requires a mechanism that allows individuals to choose tasks. The popular ‘response threshold hypothesis’ posits that some workers start engaging in particular tasks at a lower level of need than others. However, individuals may only have access to information about need after they actually engage in a task. We therefore introduce two novel interpretations of this task-allocation mechanism. While the ‘response threshold mechanism’ determines when individuals *start* working, the ‘satisfaction threshold mechanism’ drives when individuals *stop* working. We also model a ‘composite threshold mechanism’ where workers consider task need both to start and end working. Second, we model the possibility that the stimulus perceived by workers is a ‘completion’ cue instead of a ‘demand’ cue. While these may seem like subtle variations, we show here that they can yield dramatically different collective dynamics. In simulations with biologically relevant parameter ranges, response thresholds produced the quickest reaction to increases in task demand, satisfaction thresholds yielded the lowest task-switching rate, and composite thresholds most closely matched the number of workers allocated to the number needed. Different threshold types thus differentially trade off speed, cost, and accuracy. We did not model benefits of specialization; purely in terms of allocating workers to tasks, we also found that response thresholds usually perform worse than a null random choice model in terms of cost and efficiency, and variation among workers does not improve task allocation. Colonies utilizing task demand cues also tend to perform better than those using task completion cues. Our results ultimately suggest that different threshold mechanisms may be suited for different situations or types of tasks.

**Author Summary:** Division of labor is a phenomenon where workers in a community consistently differ in the tasks they work on. Many scientists believe division of labor arises in social insects (i.e. ants and bees) as a result of difference in workers’ responsiveness to cues that correspond to the demand for work in a task. For example, some ants in a colony start feeding brood much sooner than others, possibly because of a higher sensitivity, or lower ‘response threshold’, to brood pheromone. We show that instead of using such a cue to decide when to start on a task, theoretically workers may instead use it only to decide when to stop working; similarly, workers may use a cue that tells them how much work is needed in a task, or they may use one that corresponds to how much work has already been done. These seemingly subtle differences affect how much a colony invests in work and how quickly stability is reached when the balance of work needed in different tasks changes. Therefore, these different mechanisms may evolve to solve different problems.

## Introduction

Division of labor is a widespread phenomenon that increases productivity in human societies (Smith, 2010), is a defining feature of evolutionary transitions of individuality (Szathmáry & Smith, 1995), and is present in organisms as diverse as algae (Kirk, 2005), meerkats (Manser, 1999), and *E. coli* (Brameyer et al., 2022). Division of labor among social insects is usually defined as a stable pattern of variation among workers within a colony in the tasks they perform (Oster & Wilson, 1978), and is often credited for the ecological success of social insects (Wilson, 1985). Despite its near universal presence among social insects (Hölldobler & Wilson, 2009), it is unclear exactly how division of labor arises. While a number of different proximate mechanisms have been proposed to be used by social insects in allocating workers to tasks (reviewed in Beshers & Fewell, 2001; Duarte et al., 2011), one of the most widely proposed is that response thresholds are the main mechanism generating division of labor (Bossert & Wilson, 1963; Bonabeau et al., 1996; Page & Mitchell, 1998; Arcuri & Lanchier, 2017; Chen et al., 2020; Merling et al., 2020; Caminer et al., 2023). The popularity of this mechanism, in part, arises from the fact that this is a computationally simple algorithm that can achieve strong differences in specialization despite initially similar workers while simultaneously ensuring that the colony can respond to sudden increases in task demand (Page & Mitchell, 1998). These are properties that have inspired task allocation algorithms in swarm robotics (Zhang et al., 2007). However, this hypothesis is limited in its ability to predict patterns of division of labor in real social insects (Garrison et al., 2018; Leitner et al., 2019), and it does not solve all problems associated with task allocation (Jandt & Dornhaus, 2014; Berenbrik et al., 2018; Ultrich et al., 2018; Dornhaus et al., 2019; Leitner, 2019; Weidenmüller et al., 2019; Yamanaka et al., 2019; Ulrich et al., 2021). We therefore expand on the general idea of thresholds-based decision rules for task allocation to uncover new useful characteristics of this type of decision rules in any complex system, while simultaneously laying the groundwork for new interpretations of experimental data in social insects and elsewhere.

In social insect literature, ‘task allocation’ is sometimes defined as the *pattern* of how colony effort is distributed across multiple tasks (Fewell & Harrison, 2016). In the context of this paper, however, we focus on the *process* by which particular workers choose to engage in different tasks (Gordon, 1999) such as foraging, brood care, and nest construction, and call this process ‘task allocation’. The response threshold mechanism is an individual-based decision rule by which the task allocation process may occur. The response threshold hypothesis states that working social insects vary in sensitivity to stimuli that signal the need for a task to be performed (task-associated stimuli or ‘task demand cues’). Specialists in a task could be those individuals that are the most sensitive to a task demand cue (Kohlmeier et al., 2018). These individuals will perform the task first if demand increases, i.e. start their work at the lowest levels of task demand, and through their work will maintain the stimulus at a low level, i.e. below that of higher threshold workers, who never perform that task (Robinson, 1987). Thus these small internal differences might cause strong and consistent differentiation in behavior (Pankiw & Page, 2000). Even while response thresholds are fixed on the timescale of changes in task demand, they allow for dynamic changes in the number of workers allocated to specific tasks as demand for work in different tasks changes. As demand increases in one task, workers are automatically reallocated as that task’s demand cue crosses their response thresholds; if demand climbs high enough, this will happen even for high-response-threshold workers (Beshers & Fewell, 2001; Duong & Dornhaus, 2012). It is worth noting that the distribution and values of the response thresholds among workers and among tasks may have a strong effect on the colony-level outcome, and thus should maybe be considered a part of the strategy; however this has rarely been studied (Jeanson et al., 2007; Pinter-Wollman et al., 2012). A common assumption is that the response threshold values are allocated randomly across tasks and workers (Jeanson et al., 2007; Ulrich et al., 2018; Ulrich et al., 2021). An individual’s threshold values may be caused or modulated by body size, age, or differences in neural anatomy, or may be determined by stochastic processes during development (reviewed in Leitner, 2019).

Here we expand on two of the underlying components of the response threshold model: the task-associated cue and the response threshold itself. Task-associated cues are generally assumed to increase over time as the need for work increases (Bonabeau et al., 1996). For instance, in bees, brood may release more brood pheromone if left uncared for, and experimentally increasing the concentration of brood pheromone leads to an increase in pollen foragers to supplement the brood in *Apis mellifera* (Pankiw et al., 1998). Thus, the overall level of brood pheromone in the nest air may serve as a task demand cue, communicating the ‘hunger level’ of brood.

However, a task-associated cue does not necessarily need to *increase* as the need for work increases. For example, the amount of pollen stored reflects the need for more pollen foraging, and if it acts as a work cue, it is one that *decreases* as the need for work increases (because less pollen stored equals more need for pollen foraging activity). For the purposes of this paper, task-associated stimuli will be grouped into two categories: task demand cues (where the stimulus increases over time when the task is not attended to) and task completion cues (where the stimulus decreases over time when the task is not attended to).

The response threshold itself can also be inverted. There are two decision points in most formulations of the response threshold model, namely when to *start* and when to *stop* performing a task. Typically, response thresholds dictate when a worker will start a task, but workers will randomly stop performing the task (Bonabeau et al., 1996; Theralauz et al., 1998), perhaps reflecting a situation in which a worker may only be able to perceive the cue when in an inactive state (e.g. patrolling). On the other hand, if a worker can only perceive a task cue when they are currently working on the task, then a similar threshold might exist governing when a worker stops performing a task; we call these “satisfaction thresholds” (because they indicate when a worker will be ‘satisfied’ with their work). In the latter case, individuals that have a high threshold for stopping a task become specialists, as they perform the task longer than those with a low threshold. For example, Kwapich and Tschinkel (2016) found that foragers actively inhibited other ants from foraging, perhaps to signal that enough resources had been collected. Workers insensitive to this “off” cue may continue to forage regardless. These workers could have a high satisfaction threshold to that task completion cue, which could result in differences in the average time spent performing tasks across individuals (Weidenmüller, 2004; Ulrich et al., 2021).

It is then possible to create four models of the task allocation process, representing four different possible specific decision rules of individual behavior rules for choosing tasks: using either a task *completion* cue or a task *demand* cue, and either *satisfaction* or *response* thresholds. It is also possible that workers use both response and satisfaction thresholds, i.e. both the start and end points of work on a task are defined by task-cues (so that neither is completely random). We thus model two additional decision rules (one with each cue type): the composite threshold task demand model and the composite threshold task completion model. In this paper, we assume that all these thresholds are fixed and probabilistic (where the probabilities of performance are conditioned on the cue). The key assumptions and differences between these models are summarized in Table 1. We examine here the results of individual-based, stochastic simulations using each of these task allocation models, programmed in MATLAB (see Methods). In these spatially implicit models, a discrete number of workers have the choice of resting or performing different tasks, where there is a sudden increase in the demand for these tasks followed by a stable low influx of demand for all tasks. The probability that an ant chooses to start performing one of these tasks, continue to perform their current task, stop performing their task, or continue to rest depends on the strength of the demand for that task, the individual’s threshold value, and what decision rule they are utilizing. Parameters from the model are drawn from random distributions whose ranges are either measured in real ants or are assumed so that we cover a large number of possible situations.

These models are also ranked depending on how well they maximize or minimize three different metrics of task allocation. First, there is the need to dynamically adjust the number of workers engaged in the task. Doing it well without utilizing too many or too few workers can be construed as a measure of efficiency in task allocation (Dornhaus et al., 2020). Second, there are scenarios where task allocation needs to happen quickly regardless of the availability of specialists, such as with nest defense (Jandt et al., 2012; Baudier et al., 2019; Jeanson, 2019). Finally, in the design of any algorithm, accuracy and speed often trade off with cost (Marshall et al., 2006). Here, a number of costs can be envisioned; we focus our analysis on the cost of state switching (Jeanne, 1986). As state switching has been linked to temporal delays (Leighton et al., 2017) and tasks tend to be located in different parts of the nest (Sendova-Franks & Franks, 1993; Jandt et al., 2009; Guo et al., 2020; Sharma & Gadagkar, 2023), reducing state switching rates can potentially save on energetic costs.

Our goals are threefold. First, we want to determine whether or not each type of decision rule is capable of producing biologically feasible levels of division of labor as well as responding dynamically to a change in task demands. Second, we want to discover whether or not task allocation mechanisms using these different thresholds lead to different colony-level behaviors. If they do, these variations in decision rules are not entirely arbitrary, but may be under selection. In addition, the predictions from the models might help elucidate which decision rule is used by real social insects. We primarily investigate the three-way tradeoff between accuracy, speed, and cost (de Froment et al., 2014). Finally, we determine whether or not the behavior of the colony significantly changes with the number of tasks, other parameters of the model, and whether or not these threshold-based decision rules outperform a null model in which individuals are identical and do not respond directly to the cue. The probabilities of switching between states in this null model are constant across individuals and time, and are selected in such a way that the right number of workers are active every time step to complete the amount of work at hand.

## Results

### Overall performance

Each modeled decision rule (except random choice), i.e. each cue-threshold type combination, was capable of producing some level of division of labor (Fig. 1A), in the range of what is empirically seen in social insects (Gorelick et al., 2004; Dornhaus et al., 2009; Jandt et al., 2009; Holbrook et al., 2011; Holland et al., 2020; Sharma & Gadagkar, 2023), although those using ‘completion cues’ tended to be on the lower end of this range. ‘Random choice’, as expected, produced essentially no division of labor (Fig 1A). To determine whether the decision rules achieved an adaptive adjustment to task demand, we measured how much of the stimulus the colony removed from the environment (task demand) or added to the environment (task completion) relative to the best and worst possible cases (see S9 Appendix for derivation; Fig. 1B). This measure can be conceived as a measure of accuracy, as it directly measures how well the task was completed (an accuracy of 0 indicates that no work was done to resolve the cue, 1 means the cue was maximized or minimized as quickly as possible). We used timesteps to equilibrium as a measure of speed (Fig. 1C, for definition see Methods section), and the number of times workers switched their state as a measure of cost (Fig. 1D). This state switching rate includes transitions into and out of the rest state, but results are similar if we ignore these transitions and only focus on switches between tasks (S8 Appendix). The three threshold types and the random choice model differed significantly from each other in all three measures for both task demand and task completion cues (Kruskal-Wallis tests, all df = 3, all p-values < 0.001). Additionally, nearly all pairwise comparisons (between any two threshold types or random choice) were statistically significant for all response variables (all Dunn test p-values < 0.001) with only a single exception (p-value > 0.05 for composite threshold/satisfaction threshold comparison for timesteps to equilibrium). We found that the composite threshold model had the highest accuracy value, followed by satisfaction thresholds, random choice, and then response thresholds (Fig 1B). Response thresholds took the least time to reach equilibrium, followed by composite thresholds which are tied with satisfaction thresholds, and then random choice (Fig 1C). Finally, the satisfaction threshold decision rule yields the lowest cost (in terms of switching), followed by composite thresholds, random choice, and then response thresholds (Fig 1D). All of these results are comparing rules across a range of parameter values as detailed in Table 2.

### Cue type

To evaluate the effect of cue type, we performed Wilcoxon matched-pairs signed-rank tests within each threshold type and for each response variable while using the Holm-Bonferroni method to correct for multiple comparisons (p-values given in Fig 1). In many cases, cue type had a significant effect on the outcome metric if the decision rule was a threshold mechanism, but the cue type had no effect in the random choice rule (as expected, since workers don’t take the cue into account in their task choice). Task demand cues generated higher levels of division of labor in 2 out of 4 cases (Fig. 1A), higher accuracy values in 0 out of 4 cases (Fig. 1B), reached equilibrium more quickly in 0 out of 4 cases (Fig. 1C), and led to fewer state switches in 2 out of 4 cases (Fig. 1D).

**Fig 1:**
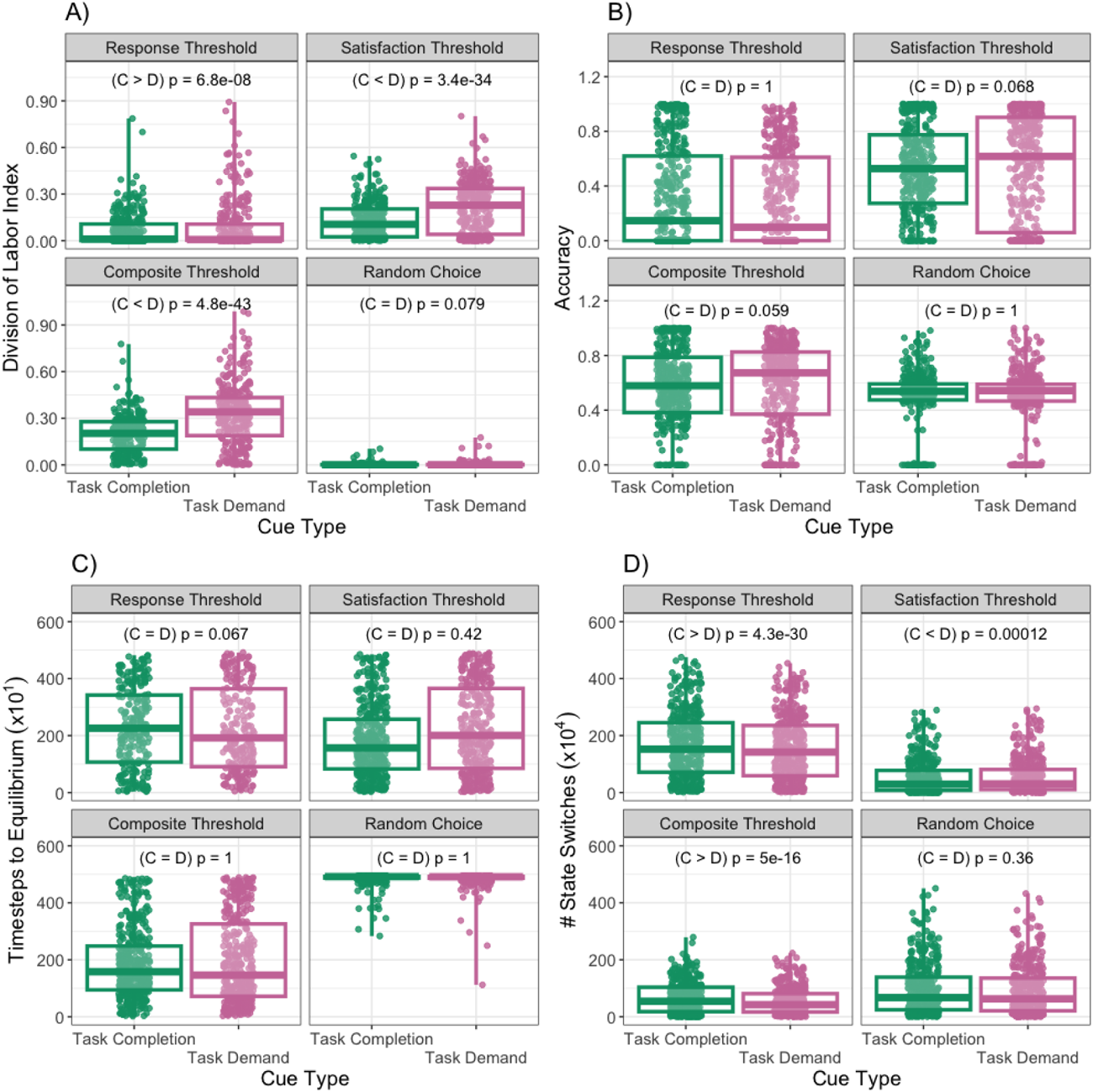
Performance metrics across threshold types. Each major panel represents a different behavioral metric while each subpanel gives a different decision rule. Boxplots show each cue type. Each point represents a single simulation, so there are differences between observations due to stochastic variation within simulations as well as differences in parameters across simulations. However, each cue type experienced the same set of parameter combinations, and p-values comparing cues are the result of Wilcoxon matched-pairs signed-rank tests. If the median of task demand is higher than that of task completion, then we denote the relationship (C < D). If it is the other way around, then (C > D). If there is no significant difference between the two groups, then (C = D). In A), task demand always produced higher levels of division of labor for all decision rules with the exception of random choice, in B) accuracy never varied across cue types for all decision rules, in C) for response thresholds, task demand had fewer timesteps to equilibrium, and in D) task demand produced fewer task switches for composite and response threshold models, more task switches for satisfaction thresholds, and had no effect on random choice.

### Variation among individuals

The most important characteristic of all the threshold-based task allocation mechanisms is that workers differ in their inherent tendency to engage in tasks. We thus wanted to analyze the effects of the amount of such variation among individuals in a colony (measured as standard deviation of response and satisfaction thresholds, parameter *σ*). We varied *σ* between 0 and 1 and performed a multiple linear regression (MLR) with *σ,* the threshold type, and their interactions were predictor variables (Fig. 2). As expected, increasing threshold variation generates higher levels of division of labor (MLR: R^2^ = 0.3694, main effects and interaction p < 0.001, but see Fig. S7.1). However, we find that this increased level of specialization is not accompanied by a reduction in the number of state switches, which either increases or decreases depending on the decision rule (MLR: R^2^ = 04121, all effects p < 0.001). *σ* may also not affect accuracy (MLR: R^2^ = 0.096, threshold type p < 0.001, *σ* effect and interaction p > 0.05), but it increases the number of timesteps to equilibrium (MLR: R^2^ = 0.3227, all effects p < 0.001). Generally, these results show that response thresholds do not generate as much division of labor as satisfaction thresholds do. They also imply that variation among individuals is not universally beneficial; while it increases division of labor, it does not influence our measure of accuracy, it has mixed effects on cost (switching), and appears to have a negative effect on speed (time to equilibrium). There may of course be benefits of specialization in terms of increased individual performance, which we do not model here.

**Fig 2:**
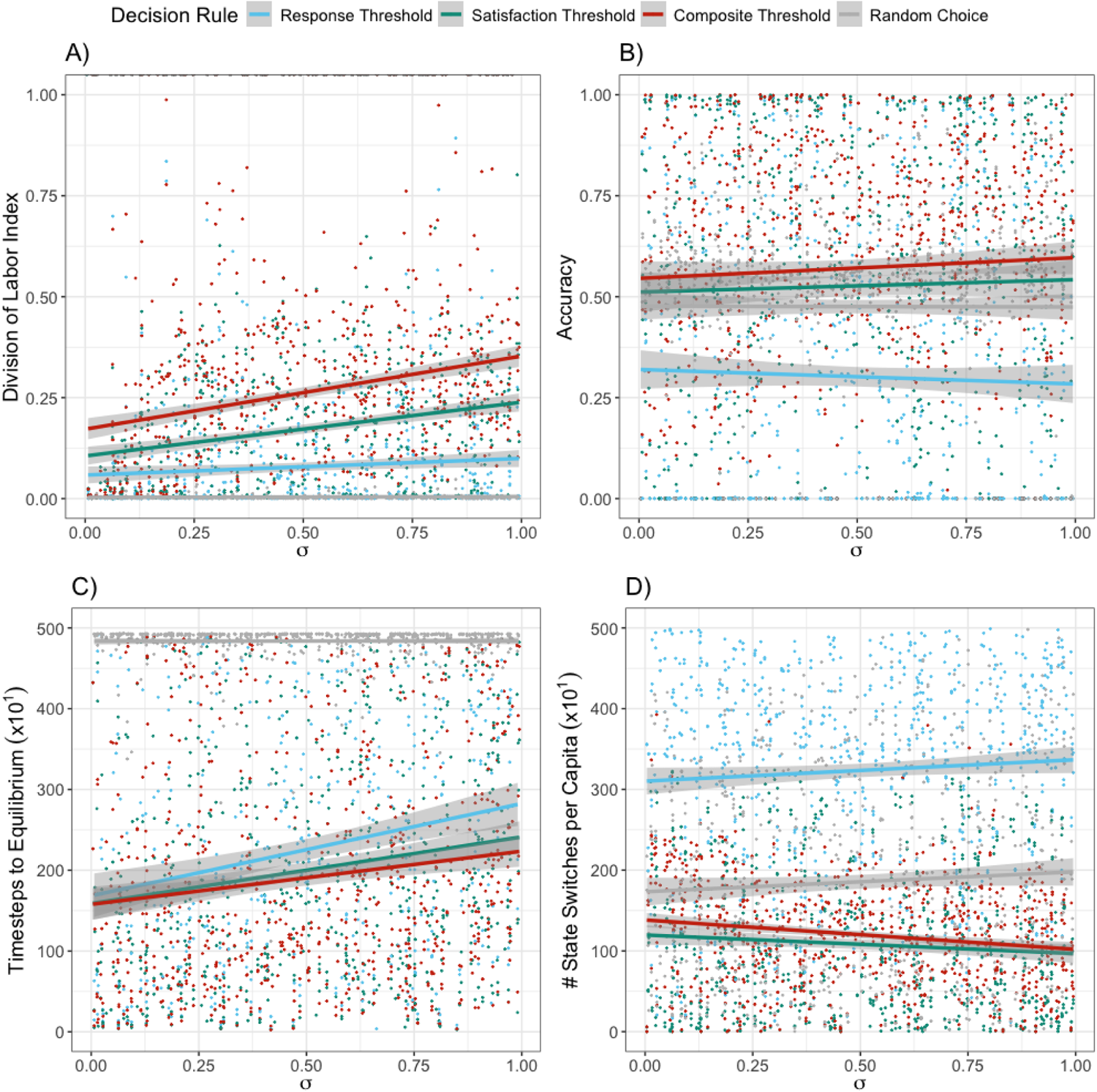
Effect of variation among workers in thresholds *σ* on performance metrics for each decision rule. Each panel represents a different behavioral metric; colors of points and lines represent the different decision rules, aggregating over both types of cues. Each point represents a single simulation (with parameter values varied as per Table 2), lines show multiple linear regressions where *σ* and the decision rule are factors. The shaded regions give standard error. Division of labor (A) depends on inherent variation among workers; we do not find an improvement in the other performance metrics with higher worker variation for any decision rule. We normalize the number of state switches (D) by dividing the total colony size *N*, as larger colonies will have more switches by virtue of having more workers. As *σ* does not influence the random choice model at all - there are no thresholds to vary - the performance of the random choice model is flat with *σ* for all performance metrics.

### Group size

We randomly varied the number of workers in each colony (*N*) between 10 and 1000 to explore the effect of group size on the four performance metrics, and to see whether the performance of different allocation mechanisms was affected by group size (Fig. 3). State switching rate (MLR: R^2^ = 0.4076, p-values for threshold type effect < 0.05, *N* and interaction effect > 0.05) and accuracy (MLR: R^2^ = 0.0972, p-values for threshold type effect < 0.001, *N* and interaction effect > 0.05) are essentially unaffected by group size, but the number of timesteps to equilibrium tends to increase with group size (MLR: R^2^ = 0.4076, p-values for *N* and threshold type effect < 0.05, interaction effect > 0.05). Division of labor also tends to decrease with colony size (MLR: R^2^ = 0.373, p-values for *N* and threshold type effect < 0.001, interaction effect > 0.05).

**Fig 3:**
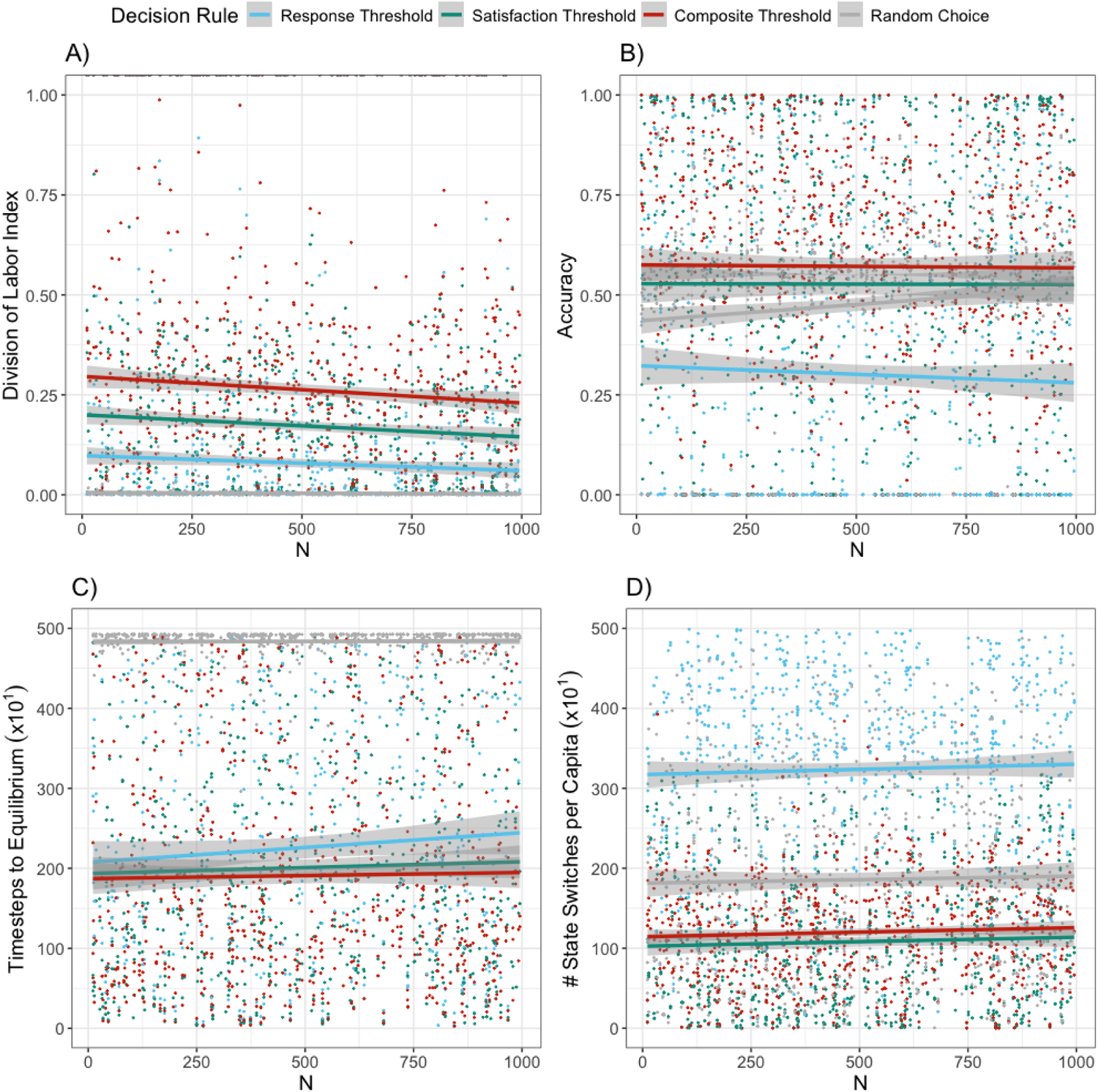
Effect of colony size *N* on performance metrics for each decision rule. Each panel represents a different behavioral metric; colors represent different decision rules; each point is a single simulation, lines show multiple linear regressions where *σ* and the decision rule are factors. The shaded regions give standard error. In our simulations, we assumed that the total amount of work scales with the number of workers (*N*, or colony size).

### Number of tasks

The relationship between the number of tasks *T* and each performance metric was evaluated individually for each decision rule with a Spearman’s rank correlation test while using the Holm-Bonferroni method to correct for multiple comparisons (p-values and correlations given in Fig 2). In general, *T* had a significant effect on some decision rule / response variable combinations, but even in these cases the effect size is small at best (Gignac & Szodorai, 2016), so the effect of *T* is negligible.

**Fig 4:**
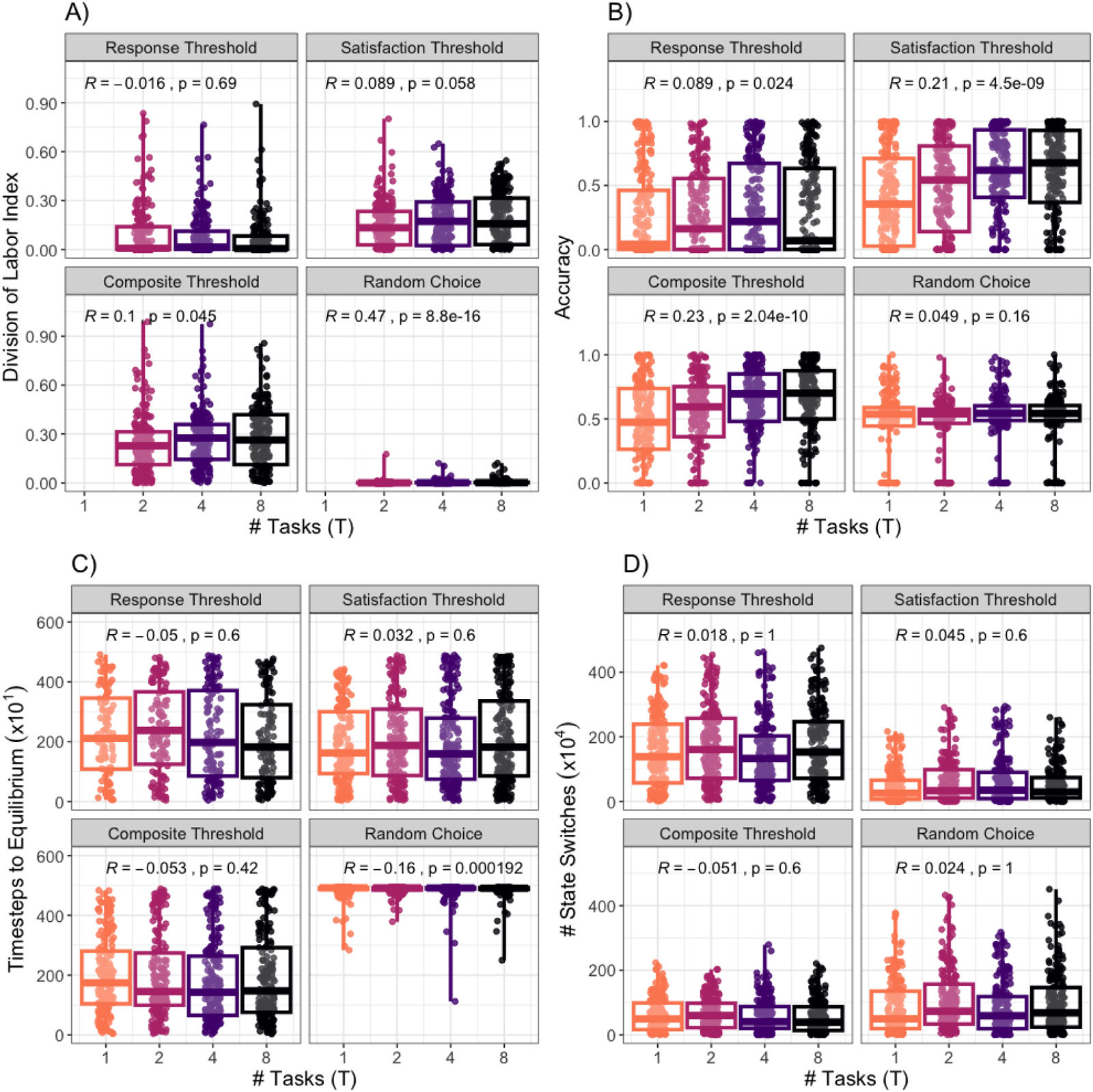
Performance metrics for differing number of tasks across decision rules. Each major panel represents a different behavioral metric while each subpanel gives a different decision rule. X-axes show the number of tasks *T*. Each point represents a single simulation, so within each boxplot there are differences between observations due to stochastic variation within simulations as well as differences in parameters across simulations. The correlation coefficients (R) and p-values are from Spearman’s correlation tests comparing each response variable to *T*. Positive values of R indicate a positive relationship between *T* and a response variable, whereas negative values indicate a negative relationship.

## Conclusion

‘Response thresholds operating on a task demand cue’ remains one of the most commonly proposed mechanisms for task allocation in social insects and for generating division of labor (Jeanson & Weidenmüller, 2014), as variation in such thresholds among individuals can both theoretically generate division of labor in virtual social insects (Bonabeau et al., 1996; Jeanson, 2007, Fontanari et al., 2023) and variations in such thresholds have been linked to division of labor in real social insect colonies (Page & Mitchell, 1998; Perez et al., 2013). Here, we show that the purported features of such a decision rule based on a task demand stimulus can also be generated by other, similar decision rules. That is, we demonstrate that satisfaction threshold decision rules, decision rules with a task completion cue, and decision rules with both response and satisfaction thresholds can each produce clear division of labor, unlike random choice models (even those correctly anticipating the needs of the colony), which do not.

Although all versions of threshold-based decision rules produce division of labor, we also demonstrate that they differ in their performance in other metrics, which may trade off with each other. This implies that different threshold decision rules might evolve for different tasks or in ecological or social situations where different aspects of performance are under selection.

Our study also examines the effects of other parameters on task allocation. In our simulations, where work demand was assumed to increase linearly with colony size, larger colony size led to a decrease in division of labor, possibly due to a lower influence of stochastic fluctuations on worker task choices. While this result may be counterintuitive given empirical work that larger colonies exhibit more division of labor (Holbrook et al., 2011; Ferguson-Gow et al., 2014), it is consistent with prior theoretical work showing that it is not task allocation mechanisms per se that generate an effect of group size on division of labor; it is only additional assumptions, like non-linear scaling in demand or efficiency, or lower relative fluctuations in task demand, that result in higher division of labor at larger group sizes (Dornhaus et al., 2012; Radeva et al., 2017).

In our simulations, each threshold type tended to maximize a different metric in the three-way trade off between efficiency, speed, and cost. First, we found that composite thresholds are the best at completing the task such that there is little residual need to perform it throughout the simulation. This accuracy could be critical for high priority tasks such as thermoregulation, where even small deviations from optimal temperatures can be extremely harmful to the hive (Becher et al., 2010; Zhao et al., 2021; Gal & Kronauer, 2022). Additionally, this high degree of performance does not come at the cost of activating too many workers for composite thresholds (S9 Appendix), which can also be critical for thermoregulation, as it is an energetically costly set of tasks (Bretzlaff et al., 2023). In other cases, the speed of a response may be more important, such as in nest defense (Robinson, 1987; Jongepier & Foitzik, 2016). We found that colonies that employed response thresholds were the quickest to reach equilibrium after the initial shock of stimulus, indicating that this type of threshold produced faster-responding colonies than satisfaction or composite thresholds. Conversely, when task switching is particularly costly (e.g. because of the importance of learning or physiological specialization), the task switching rate may be the most relevant performance metric. We find that satisfaction thresholds minimize task-switching relative to the other tasks, and so could be used to regulate tasks which have a high barrier to entry (Holway, 1998), are spatially distant from other tasks (Mersch et al., 2013), or can cross-contaminate other tasks (Casillas-Pérez et al., 2023). For instance, when undertakers were removed from a honey bee hive, they were not soon replaced with other workers (Robinson & Page, 1995). This violates predictions of the response threshold hypothesis, but is not necessarily an issue for the satisfaction threshold hypothesis, as even workers with a low satisfaction threshold (who would not stop undertaking until the oleic acid was removed from the environment) may not start the task for a long period of time.

In addition to finding strong differences in performance across threshold types, we also found differences between decision rules which used either task demand or task completion cues, although there was no effect of cue type in the random choice model. This suggests an interaction between cue type and decision rule. Generally, we find that task demand cues cause colonies to have higher levels of division of labor, have fewer timesteps to equilibrium, and switch tasks less often, although there are exceptions to this rule for some decision rule - cue type combinations. In many circumstances, then, it may be better to regulate a task with a demand cue compared to a completion cue, but this may not always be possible depending on the nature of the task at hand. Given the lack of an effect with the random choice model though, this distinction between cues may not matter for other types of non-threshold-based decision rules.

In summary, the often-cited verbal hypothesis of task allocation by response to task-associated cues can in fact be interpreted to imply that workers respond to such cues when starting *or* stopping tasks, and that cues might reflect task demand or task completion, or some combination of these. Here we show that these details of a task allocation mechanism can have important consequences for collective performance, and in particular, that different performance metrics may trade off against each other. Indeed, we found that the traditional response threshold hypothesis performed poorly relative to other decision rules, as it organized labor in such a way that it was not accurate, workers switched tasks often, and it tended to activate fewer workers than what was necessary (Appendix 9), perhaps illustrating the difficulty of resolving response thresholds empirically. These insights will be important in understanding the evolution of task allocation in biological systems, the design of appropriate experiments to test hypotheses about task allocation mechanisms, and in optimizing task allocation and system-level outcomes in engineered systems such as in distributed computing and traffic management (Castello et al., 2016; Jiang et al., 2020; Wu et al., 2020).

## Methods

### Decision rules for a single task

We begin by discussing our discrete time model in the context of one task, before moving onto the multi-task case. In this simple case, we treat each worker as an individual Markov chain with two states: an active and an inactive state. In response threshold decision rule, the probability of switching from the inactive state to the active state depends on the strength of the task associated cue, whereas the probability of leaving the active state is constant. The opposite is true for the satisfaction threshold decision rule. Both leaving and entering the active state depends on the cue with the composite threshold decision rule. When the worker is in the active state, she will either reduce the amount of the cue in the environment (task demand) or increase it (task completion). If she stays in the inactive state, the cue will increase or decrease (respectively) on its own until it reaches some maximal or minimal value.

The traditional fixed response threshold task demand rule posits that as the need to do labor increases, the probability that any workers will start working on the task will also increase. Workers with a lower response threshold will have a higher probability of starting a task than workers with a higher threshold at intermediate values for the task stimulus. Bonabeau et al. (1996) satisfied this statement with the following assumed Hill function, which generates an s-shaped curve:

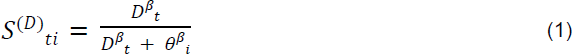

where *S^(D)^_ti_* is the probability of starting a task for individual *i* at time *t* (Fig 5) for a given value of the task demand cue *D_t_*. This variable *D_t_* represents the strength of task demand cue at time *t*, and 0 ≤ *D_t_* ≤ 1. A value of 0 indicates that the task does not need to be performed, whereas 1 indicates maximal need. As *D_t_* increases, so too must the probability of starting the task, so equation (1) generates a positively-sloped sigmoid curve where the fixed response threshold for individual *i* where *θ_i_* is the inflection point and *ꞵ* determines the steepness of the function. Both of these free parameters are constant throughout the course of a simulation. Additionally, *i* = 1, 2, …, *N* and *t* = 0, 1, 2, …, *M*.

Task completion cues, *C_t_*, have the opposite interpretation as task demand cues, where 0 represents the case of maximal need and 1 indicates completion of the task. Therefore, the slope of the resulting response curve, *S^(C)^_ti_*, should have a negative slope and it should take on the same form as equation 1. Such a function can be found by finding the complement to equation (1), substituting *D_t_* with *C_t_* :

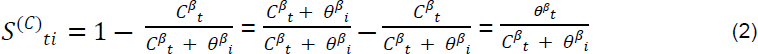

For both decision rules, the probability of leaving the inactive state (or stopping the task) is *l^(D)^_ti_*, *l^(C)^_ti_* = *p*, a free parameter of the model.

We can also use a similar set of equations for the satisfaction threshold decision rule. This rule posits that as the task demand cue increases, the probability that any workers will stop working on the task will decrease. As with equation 2, this can be accomplished with the complement to the traditional Hill function:

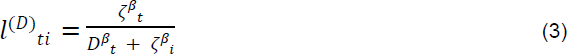

where *□_i_* is the satisfaction threshold for worker *i*. Finally, the satisfaction threshold decision rule for task completion stimuli is:

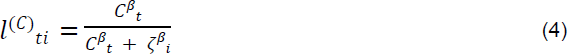

For each of these functions, *S^(D)^_ti_*, *S^(C)^_ti_* = *p*. In the case of composite threshold decision rules, there is no constant probability of either starting or stopping the task. The probability of starting a task for a task demand stimulus is given by equation 1 while the probability of stopping is given by equation 3. Conversely, the starting probability for task completion is equation 2 while the stopping probability is equation 4 (Fig 5).

**Fig 5:**
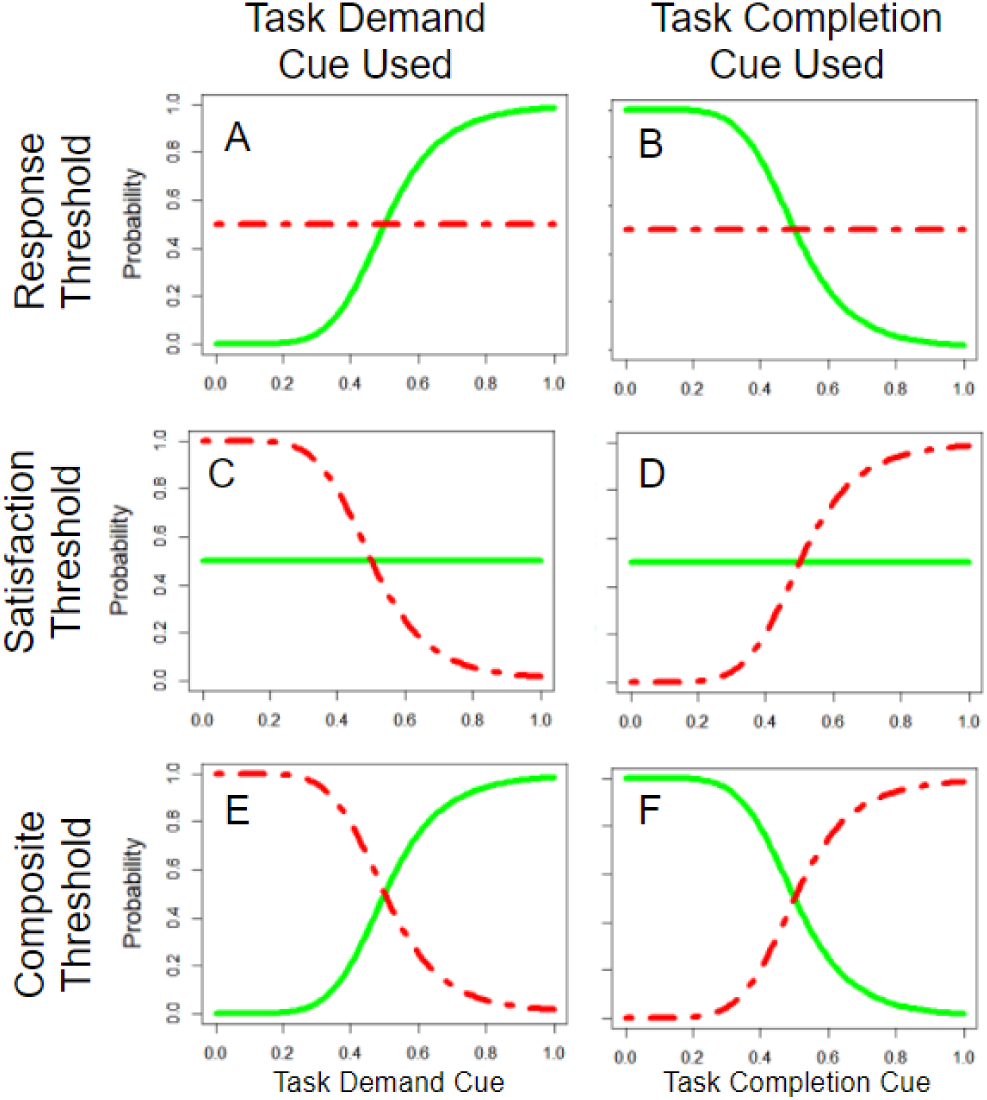
Probability of starting and stopping for each model across cue values. For all 6 possible decision rules modeled here (i.e. all combinations of using a task demand or task completion cue, and using either response, satisfaction, or composite thresholds), we show here the probability of starting (green, solid lines) and stopping (red, dashed lines) a task as a function of the task cue. The model used in Bonabeau et al. (1996) is represented in panel A: response thresholds using a task demand cue. For this illustration, we used the numerical values *□* and *□* = 0.5, *β* = 6, and *p* = 0.5.

### Update functions for task associated cues

The task demand cue in our model evolves over discrete time in much the same way as it does in the classical model (Bonabeau et al., 1996). The cue increases linearly over time, and the amount of work each worker can complete is inversely proportional to the number of workers in the colony *N*. The underlying assumption is that work scales linearly with colony size (Cole, 1986; Bonabeau et al.,1996), which may or may not be the case (Fewell & Harrison, 2016). Nonetheless, this represents a simple starting point on which we can compare different threshold decision rules. We also assume that all workers are equally efficient at any task, since we are not focusing on examining the benefits of specialization. Empirically, specialization may not influence efficiency (Dornhaus, 2008; Cole, 2020) and the ant species that we use to parameterize these models (the harvester ant *Pogonomyrmex californicus*) are monomorphic (S1 Appendix). Equation 5 satisfies these conditions:

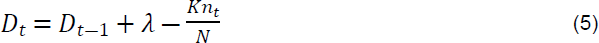

where *λ* is the growth rate of the task, *K* is the amount of the cue a worker can remove in a single timestep, and *n_t_* is the number of ants currently performing the task. In simulations, if *D_t_* < 0, then it is set to 0. Likewise, if *D_t_* > 1, then it is set to 1.

Reverting this equation to one that governs the task completion cue *C_t_* is trivial, as it only requires changing signs:

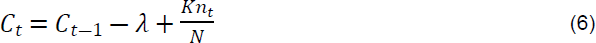

### Decision rules for multiple tasks

We now extend the model to multiple tasks. To make the performance of the task allocation rules comparable, we need to (1) modify the number of states that a worker can be in, (2) normalize the choice probabilities so that their sum always equals 1, and (3) normalize the work done so that in every model the sum of work increase across all tasks is the same, regardless of how many tasks there are.

Let *j* be the index for tasks so *j* = 0, 1, 2, …, *T*. Each ant can therefore be in *T*+1 states, where state 0 is the inactive state (Fig 6). In this model, we disallow transitions between states. A worker must stop performing a task before they can start performing a different task. We do this to remove ambiguity in the model, as it is not clear whether a direct transition from one task to another is the ant starting a new task or stopping the old task. This constraint means that when an ant is currently performing the *j*’th task, there are only two possible outcomes. The ant can continue performing the task, or she can stop. The stopping equations (equations 3 and 4) are guaranteed to be greater than or equal to 0 and less than 1, and the parameter *p* is randomly drawn from a distribution bounded between 0 and 1, there is no need for an additional correction factor. However, when the ant is inactive, then there are *T*+1 potential outcomes, so such a correction is needed. We make this correction with a new term called the standardization coefficient, which is *b^(D)^_tj_* for task demand cues and *b^(C)^_tj_* for task completion.

**Fig 6:**
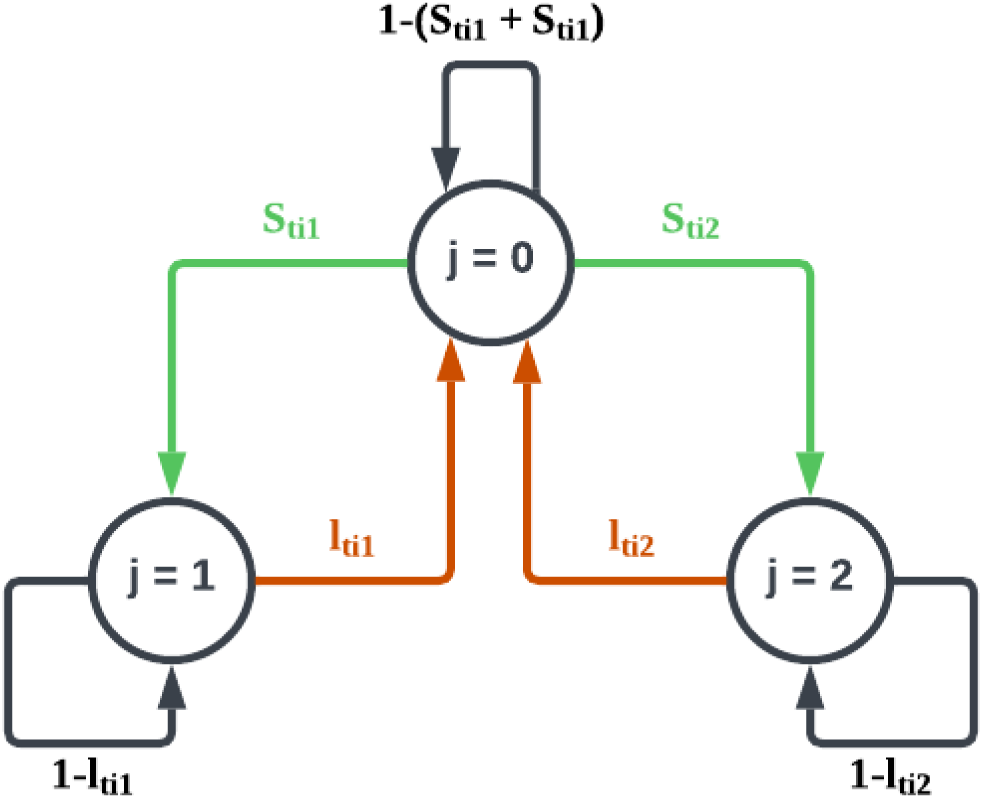
Markov chain representing transition probabilities for each worker. In the case where *T* = 2, there are three possible states that ant *i* can be in at any time *t*. When *j* = 0, the ant is in the inactive state, when *j* = 1 she is performing task 1, and when *j* = 2 she is performing task 2. We display transition probabilities generically here, ignoring superscripts associated with different task cues. The probability that she starts performing task j, then, is *S_tij_*, (green arrows), while the stopping probability is *l_tij_* (red arrows). The probability that she stays in her current state is the complement of these probabilities (black arrows).

This standardization coefficient is the relative weight attributed to task *j* when compared to all other tasks. A task with a high standardization coefficient needs to be performed more urgently than a task with a low standardization coefficient. Each standardization coefficient can take on a value between 0 and 1, and the sum of all standardization coefficients is 1. By multiplying a standardization coefficient *b^(D)^_tj_* against its associated probability *S^(D)^_tij_* (note the extra subscript *j* for task; this procedure also works for *b^(C)^_tj_* and *S^(C)^_tij_*), then the sum of probabilities for ant *i* at time *t* is guaranteed to be less than or equal to 1. This works regardless of whether or not *S^(D)^_tij_* is computed with a decision rule or is set equal to a constant. This highlights an additional assumption of the model: when workers are inactive, they are privy to the needs of all tasks, but when they are active they only pay attention to the need of the focal task. This appears to be at least partially true, as harvester ants are less likely to respond to alarm pheromone when they are already engaged in brood care (Guo, 2021).

For task demand cues, this standardization coefficient is calculated as:

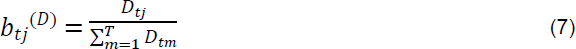

Whereas for task completion, the standardization coefficient will be the complement of the cue:

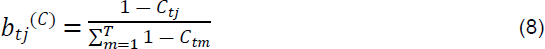

The complete set of starting and stopping probabilities which incorporate standardization coefficient for each model are given in Table 1.

To ensure that there is the same amount of ‘work’ in the environment regardless of how many tasks there are, we simply multiply an individual’s efficiency by *T* in the update formulae:

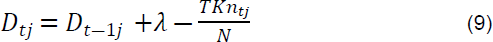

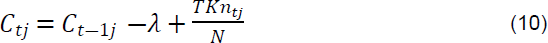

**Table 1:**
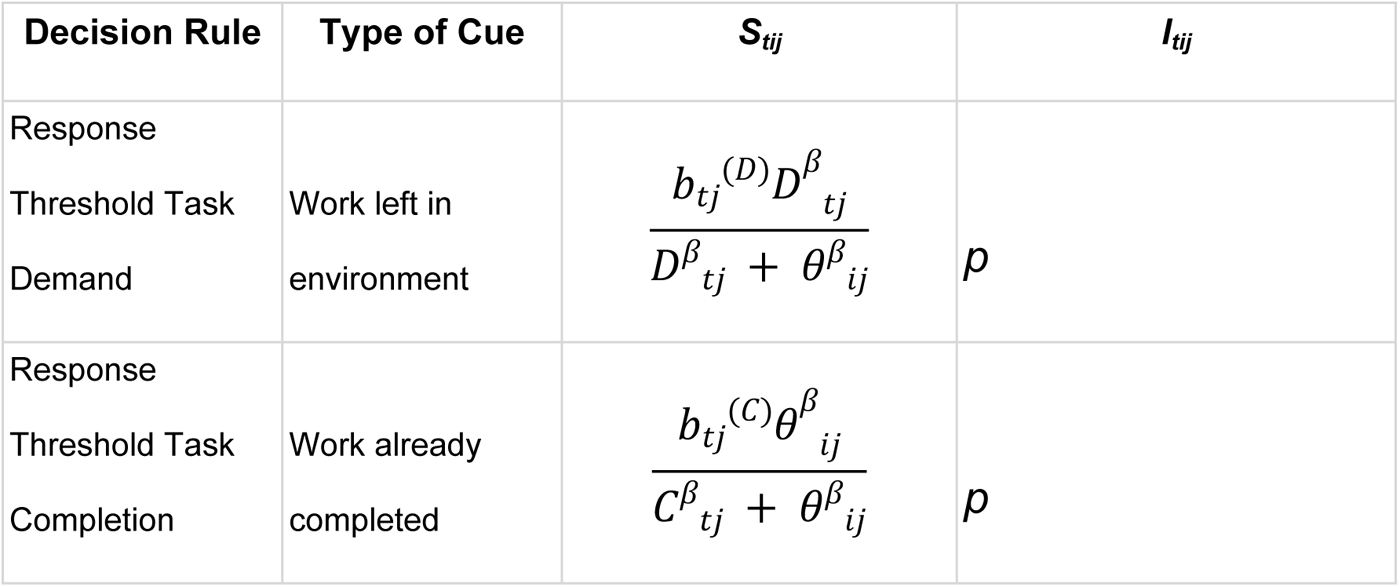
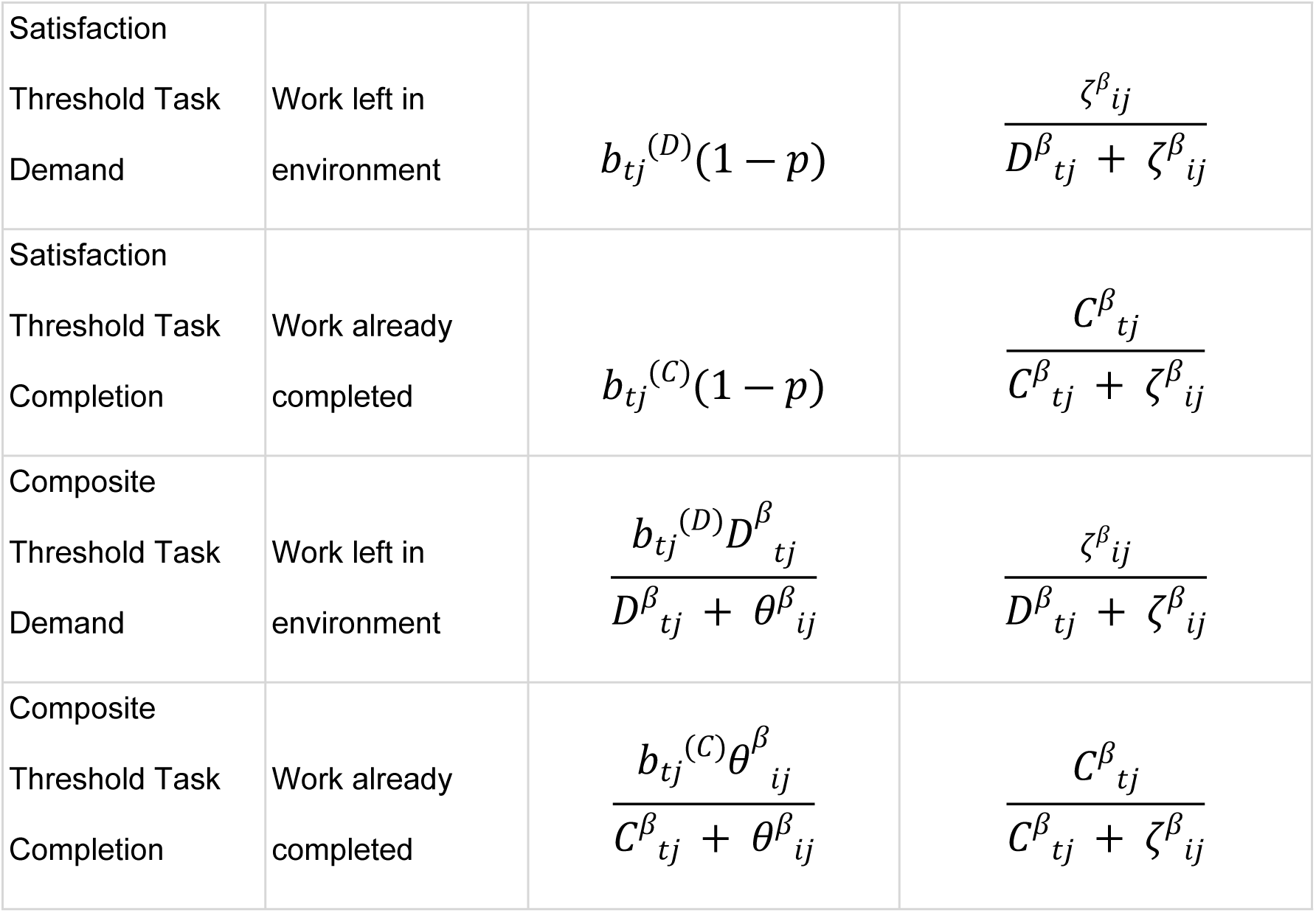
Summary of equations for each threshold model. The *S_tij_* column gives the probability that an ant will switch from an idle state to doing task j. The *l_tij_* column gives the probability of switching from a state of doing task j to the idle state.

| Decision Rule | Type of Cue | $S_{ij}$ | $I_{ij}$ |
| --- | --- | --- | --- |
| Response<br>Threshold Task<br>Demand | Work left in<br>environment | $\frac{b_{tj}^{(D)} D_{tj}^\beta}{D_{tj}^\beta + \theta_{ij}^\beta}$ | $p$ |
| Response<br>Threshold Task<br>Completion | Work already<br>completed | $\frac{b_{tj}^{(C)} \theta_{ij}^\beta}{C_{tj}^\beta + \theta_{ij}^\beta}$ | $p$ |
| Satisfaction<br>Threshold Task<br>Demand | Work left in<br>environment | $b_{tj}^{(D)}(1 - p)$ | $\frac{\zeta_{ij}^\beta}{D_{tj}^\beta + \zeta_{ij}^\beta}$ |
| Satisfaction<br>Threshold Task<br>Completion | Work already<br>completed | $b_{tj}^{(C)}(1 - p)$ | $\frac{C_{tj}^\beta}{C_{tj}^\beta + \zeta_{ij}^\beta}$ |
| Composite<br>Threshold Task<br>Demand | Work left in<br>environment | $\frac{b_{tj}^{(D)}D_{tj}^\beta}{D_{tj}^\beta + \theta_{ij}^\beta}$ | $\frac{\zeta_{ij}^\beta}{D_{tj}^\beta + \zeta_{ij}^\beta}$ |
| Composite<br>Threshold Task<br>Completion | Work already<br>completed | $\frac{b_{tj}^{(C)}\theta_{ij}^\beta}{C_{tj}^\beta + \theta_{ij}^\beta}$ | $\frac{C_{tj}^\beta}{C_{tj}^\beta + \zeta_{ij}^\beta}$ |

### Simulations and metrics of performance

We simulated the models on a simple cue shock response task, as follows. Colonies of size *N* were generated by drawing threshold values for each ant from a truncated normal distribution (range is between 0 and 1) whose variance is a free parameter (*σ*) and whose mean is *μ*. The thresholds for an individual were independently drawn for each task (the ‘idiosyncrasy’ case in Pinter-Wollman et al., 2012, which was mostly empirically supported for disparate tasks). Individuals with composite thresholds had independently drawn response and satisfaction thresholds. As these thresholds were independent of one another, the starting threshold was not necessarily lower than the ending threshold, meaning that for high values of *β,* a worker that switches onto a task would immediately stop it. However, at low levels of *β,* this outcome is not guaranteed so she may continue working for a few timesteps.

Once these colonies were generated, they were subjected to a sudden “shock” of task demand (i.e. an abrupt, one-time increase in the task demand cue or decrease in the task completion cue): in simulations with a task demand cue, the initial cue strength for all tasks is 1 (*D_0j_* = 1 ∀ *j*), while for task completion cues the initial cue strength is 0 (*C_0j_* = 0 ∀ *j*). After this shock occurs, *D_tj_* and *C_tj_* change over time according to equations 9 and 10, depending on how workers behave. All workers start in the inactive state, so the number of workers performing each task is initially 0 (*n_0j_* = 0 ∀ *j*). Workers in the colony then switch between the inactive state and different tasks according to the probabilities listed in Table 1. The simulation then runs for 5,000 timesteps before it terminates. *N*, *p*, *K*, *β*, *μ* and *σ* are free parameters of the model (Table 2). At the onset of each simulation, values for these parameters are randomly selected from either a discrete uniform distribution (*N*), a continuous uniform distribution (*p*, *β*, *μ* and *σ*) or an inverse uniform distribution, which is the distribution which results from dividing 1 by a uniform random variable (*K*; S6.1 Appendix). We aimed to cover a biologically plausible range within the allowed parameter ranges. However, many of the parameters are difficult to estimate from the empirical literature. We used data collected from 25 *Pogonomyrmex californicus* colonies to try to estimate some values for *N*, *p*, and *K* (S1 Appendix). *λ*, the change of the cue intensity per timestep, is held constant, as the impact of this parameter depends on its relative size to *K*, so only *K* needs to be varied. Free parameters that represent different aspects of the thresholds (*β*, *μ* and *σ*) are part of the task allocation mechanism. Empirical estimates are essentially absent, and we aimed to cover a broad range (S6.1 Appendix). The effects of each parameter on simulation outcomes were evaluated with a sensitivity analysis using partial correlations (S3 Appendix).

**Table 2:** Description for how parameter values were selected for each simulation. Here we give the descriptions of each parameter, the upper and lower limits for each parameter, and a qualitative description of how the parameters were chosen (S1 Appendix and S6.1 Appendix).

| Parameter Name | Parameter Range | Parameter Selection |
| --- | --- | --- |
| Number of Tasks: $T$ | {1, 2, 4, 8} | Iterated through each task number |
| Worker efficiency (the amount of work a single worker can perform in a single timestep): $K$ | [0.001, 0.1] | Drawn from continuous inverse uniform distribution |
| Colony Threshold Mean: $\mu$ | [0, 1] | Drawn from continuous uniform distribution |
| Colony Threshold Standard Deviation: $\sigma$ | [0, 1] | Drawn from continuous uniform distribution |
| Colony Size: $N$ | [10, 1,000] | Drawn from discrete uniform distribution |
| Probability of Stopping (in models with 'response thresholds') or Probability of Continuing to Rest (in models with 'satisfaction thresholds'): $p$ | [0.5, 1] | Drawn from continuous uniform distribution |
| Probability Curve Steepness: $\beta$ | [1, 10] | Drawn from continuous uniform distribution |
| Rate of Change for Stimulus: $\lambda$ | 0.001 | Held constant |

At the end of each simulation, we record various colony-level metrics of task performance (Beshers, 2023). We record the division of labor index for the colony (Gorelick et al., 2004; but see correction in Dornhaus et al., 2009), the total number of state switches performed across the simulation and across all workers of the colony, the number of timesteps it took for the task to reach an equilibrium, and accuracy. To find the number of state switches, we counted the number of times workers switched from any state, both into and out of the rest state. We consider this measurement to represent a cost on the system, as state switching can be energetically inefficient (Goldsby et al., 2012; Jeanson & Lachaud, 2015) or may risk cross contamination of waste-borne pathogens between tasks (Hart & Ratnieks, 2002).

To calculate ‘speed’ of appropriate task allocation, we counted the number of timesteps to ‘equilibrium’. We used an operational definition of ‘equilibrium’ that, in preliminary exploration, seemed to approximate the steady-state: we found the first timestep where the cue was within 0.5% of the mean of the final 100 timesteps of the simulation. We consider this a measurement of speed as it measures how quickly the colony is able to eliminate the initial bout of task need (stimulus).

Accuracy is our measure of the quality of task allocation that captures how much work the colony did relative to how much was necessary to deplete the task demand cue (or maximize the task completion cue; Dornhaus et al., 2019) over the course of the simulation. This value is scaled such that an accuracy value of 0 reflects a situation in which workers are never active, and thus the cue is maximally high (task demand) or low (task completion) over the course of the simulation. A value of 1 indicates that the cue is as low as possible (task demand) or as high as possible (task completion) given constraints on worker efficiency (S9 Appendix). This can be accomplished by all workers performing work every single timestep, but this is not necessarily a positive outcome. High accuracy values can still be accomplished even with many workers resting (S4 Appendix, S9 Appendix). We consider this value to be a measure of accuracy as it captures the quality of the work performed: a value of 1 indicates that there is no residual work leftover.

In order to not only compare these decision rules to one another, but also to test whether their basic tenets - variation among individuals and responsiveness to global information about g the need for work - contribute to colony performance, we also introduce a ‘null model’ as a baseline. In this null model, we assume that all workers in a colony have an equal and constant probability of entering and leaving different tasks, irrespective of the task stimulus (S5 Appendix). We call this decision rule ‘random choice.’

We ran 100 simulations for each combination of threshold types and cue type (the time course of example simulations is given in Figs 7 & Fig S6.1). There are 4 decision rules, 2 cue types = 8 combinations, and as there is also a set of 4 numbers of tasks that we iterate through, there are 3,200 simulations total. Across the 100 replicate runs for each of these combinations, we randomly varied the other parameters (according to Table 2). In the random choice model, the parameters *β*, *μ*, and *σ* do not apply as there are no thresholds. There is just the probability of starting any task, *S*, as well as the probability of leaving a task, *L*, both of which are constant across timesteps, tasks, and individuals. Due to the constraints of the model, these two parameters cannot be varied independently of one another. While they are randomly drawn for each simulation, many combinations of values that do not meet a certain condition are rejected such that the final value ensures that, on average, enough workers are activated to remove the continuous inflow of the cue as well as the initial shock of task demand (Fig 7; S5 Appendix). All simulations were carried out using MATLAB and Statistics Toolbox Release 2022b (The Mathworks Inc., 2022).

**Fig 7:**
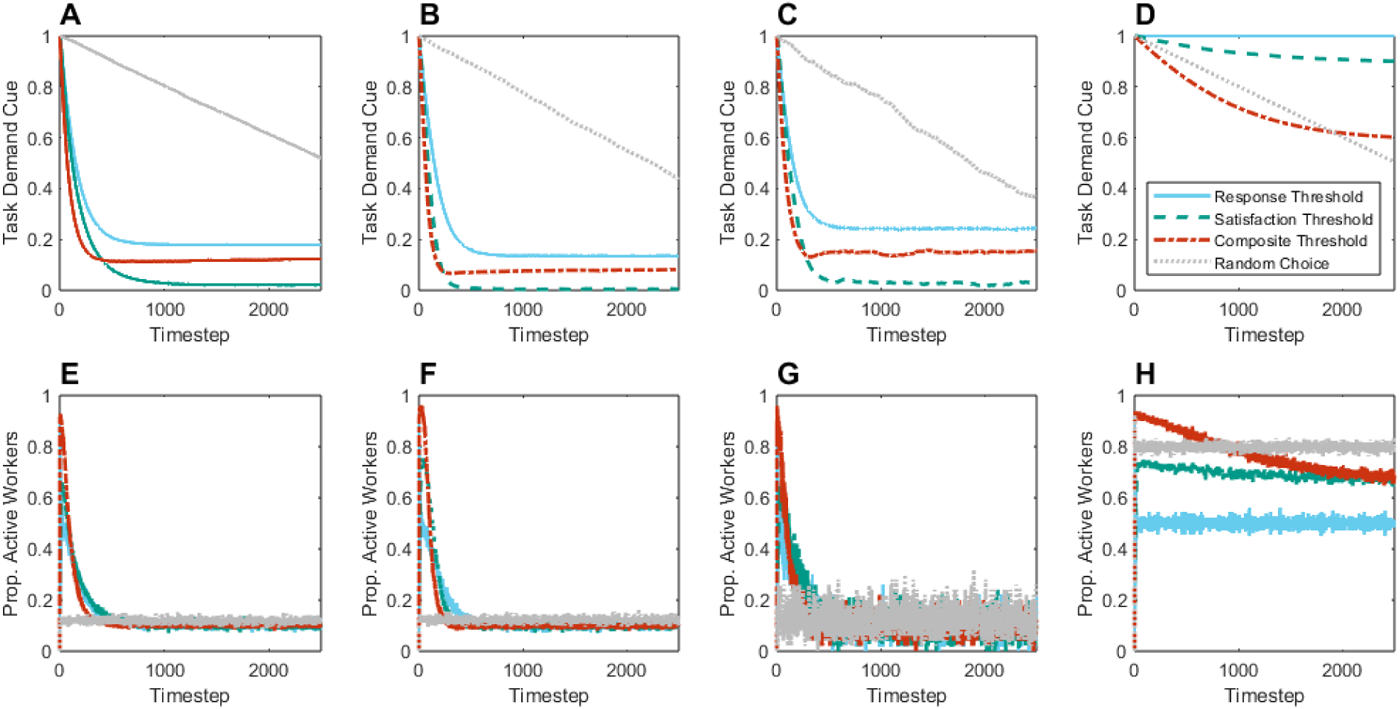
Behavior of the model over the course of a simulation run using task demand cues for different parameter combinations. We illustrate here what happens over the time course of one example simulation run for each rule and several parameter combinations. The first row shows how the task demand cue changed over time for each decision rule (color) while the second row shows how the proportion of active ants changed over time. Columns correspond to different parameter combinations. For panels A/E, C/G, and D/H, there is only a single task (*T =* 1) so the task demand cue corresponds to that single task. For B/F, there are four tasks (*T =* 4), so in B the given trace is the mean demand cue across these four tasks. For A/E, B/F, and D/H, the colony size (*N*) is 1,000, whereas in C/G *N* = 100. Finally, in A/E, B/F, and C/G, the amount of work that a single ant can complete in a single timestep (*K*) is 0.01, whereas in D/H, *K* = 0.0015, illustrating a situation where the colony can barely keep up with the amount of work that needs to be completed. In all panels, *p* = 0.9, *β* = 6, *μ* = 0.5, and *σ* = 0.5. All simulations start with a high number of workers being recruited to work in response to the task demand, and then settle to a steady-state with the exception of the random choice model, which is designed to remove the initial shock of cue by the end of the simulation, so here the cue steadily decreases over time. Only the first 2500 timesteps are shown, but simulations always run for 5000 timesteps.

### Statistical analysis

We are primarily interested in characterizing the tradeoff between accuracy (the capacity to minimize residual work), speed (the number of timesteps to equilibrium), and cost (the number of state switches) between the different models. As each of these performance metrics are not normally distributed (Shapiro-Wilks test p-value < 0.01 for each metric), and transforming these variables using the Box-Cox method produced ill-fitting linear models (as determined from examining residual plots), we opted to perform nonparametric tests instead to determine the changes in these performance metrics across different model features. To see if there are differences across threshold types, we perform a Kruskal-Wallis test with a post-hoc Dunn test to determine significance groups.

To determine the effect of cue type, we performed paired Wilcoxon matched-pairs signed-rank tests between the two cue types for each response variable within each decision rule. A paired test was possible as the same set of free parameter values was run for each cue. Similarly, we tested for the effect of the number of tasks *T* by performing Spearman’s rank correlation tests between *T* and each performance metric within each decision rule.

To measure the effect of threshold variance *σ* and colony size *N* on the different performance metrics across threshold types (response threshold, satisfaction threshold, composite threshold, random choice),, we performed a multiple linear regression where the response variable was one of the performance metrics, and the predictor variables were either *σ* or *N*, the threshold type and the interaction between these two variables.

All tests were performed in R and R Studio (v4.3.1; R Core Team 2023) with the tidyverse (v4.43; Wickham et al., 2019) and dunn.test (v1.3.5; Dinno, 2017) packages.

## Acknowledgements

We thank the members of the Dornhaus, Fewell, and NRD labs for their support, Bobby Jensen for help optimizing code, and in particular Dr. Nicole Leitner for her comments on the manuscript. This work was funded by NSF (grant # DGE-1143953 and IOS-14455983).

